# Wake respirometry may quantify stress and energetics of free-living animals

**DOI:** 10.1101/2022.04.11.487318

**Authors:** Kayleigh Rose, Rory P Wilson, Claudia Ramenda, Hermina Robotka, Martin Wikelski, Emily L C Shepard

## Abstract

Quantifying activity-specific energy expenditure in free-living animals is a major challenge as current methods require calibration in the lab and animal capture. We propose “wake respirometry”, a new method of quantifying fine-scale changes in CO_2_ production in unrestrained animals, using a non-dispersive infrared CO_2_ sensor positioned downwind of the animal i.e. in its wake. We parameterise the dispersion of CO_2_ in a wake using known CO_2_ concentrations, flow rates and wind speeds. Tests with three bird species in a wind tunnel demonstrated that the system can resolve breath-by-breath changes in CO_2_ concentration, with clear exhalation signatures increasing in period and integral with body size. Changes in physiological state were detectable following handling, flight and exposure to a perceived threat. We discuss the potential of wake respirometry to quantify stress and respiratory patterns in wild animals and estimate activity-specific metabolic rates through the full integration of CO_2_ production across the wake.

## INTRODUCTION

Determination of the energy expenditure of free-living animals is pivotal for understanding the costs and rewards of behaviours and elucidating strategies that enhance lifetime reproductive success (Lemon 1991; Shaffer, Costa & Weimerskirch 2003). However, quantification of activity-specific metabolic rate in freely moving individuals is a major challenge (Wilson & Culik 1993; Butler *et al*. 2004). Methods that have provided key insight in this regard include ‘heart rate’ (Nolet *et al*. 1992; Bevan *et al*. 1994; Bevan *et al*. 1995; Ward *et al*. 2002; Green 2011) and ‘dynamic body acceleration’ as proxies for energy expenditure (Wilson *et al*. 2006; Halsey *et al*. 2009; Halsey, Shepard & Wilson 2011; Wilson *et al*. 2020). Heart rate loggers can inform us of relative costs in animals both at rest and during activity, while dynamic body acceleration allows the interrogation of costs associated with movement. However, each of these methods has to be calibrated by indirect calorimetry (Halsey *et al*. 2009; Gleiss, Wilson & Shepard 2011; Halsey, Shepard & Wilson 2011; Halsey & Bryce 2021) or doubly labelled water (Nolet *et al*. 1992; Bevan, Speakman & Butler 1995; Elliott *et al*. 2013). Both indirect calorimetry and doubly labelled water work by assessing the rate of CO_2_ production (and, in indirect calorimetry, sometimes O_2_ consumption (Lighton 2019)). A limitation of indirect calorimetry is that it necessitates animals to be confined to boxes (e.g. (Hawkins, Butler & Speakman 2000)) or equipped with masks (Ward *et al*. 2001; Morris, Nelson & Askew 2010; Langman *et al*. 2012), which prohibits or constrains expression of many behaviours and can induce stress in the study animal (Tucker 1972). In contrast, doubly labelled water allows animals to move freely within their natural environment, but does not allow easy assessment of activity-specific metabolic rate (Nagy, Siegfried & Wilson 1984; Wilson & Culik 1993) although judicious use of sophisticated animal-attached tags is providing a way forward (Sutton *et al*. 2021)).

During the years that these processes have been developed and refined, our ability to determine CO_2_ concentration with high accuracy, even at low concentrations, has advanced dramatically. In particular, non-dispersive infrared spectroscopy (NoDIS) has been shown to resolve CO_2_ concentrations as low as 0.01 ppm (https://www.licor.com/env/products/gas_analysis/LI-7000/specifications.html). Exhaling (air-breathing) animals have CO_2_ concentrations around 4% when their respiratory gases leave their body, with this concentration being diluted with distance from the source due to diffusion and wind. Nonetheless, the accuracy of NoDIS means that the CO_2_ signal should be detectable at some distance from the CO_2_ source using a system that does not interact with the study animal in any physical way. We propose that it might be possible to determine metabolic rate by positioning these NoDIS sensors close to unrestrained, and possibly even free-living, animals and examining CO_2_ concentrations over time, providing there is directional flow of air. This would necessitate mapping out a full 2D cross-section of the CO_2_ wake. We term this approach ‘wake respirometry’ because the CO_2_ signal from an animal is drifted downwind and over the sensor. The concentration of CO_2_ at any point around an animal, and therefore the ability of a NoDIS to quantify it, will depend on the concentration of CO_2_ being emitted in the exhaled air, the position of the sensor relative to the source, and the speed and direction of the wind passing over the animal (Figure 1), as well as the expiratory flow rate, and the rate of diffusion and dilution of CO_2_ in air and water vapour.

**Figure 1.**
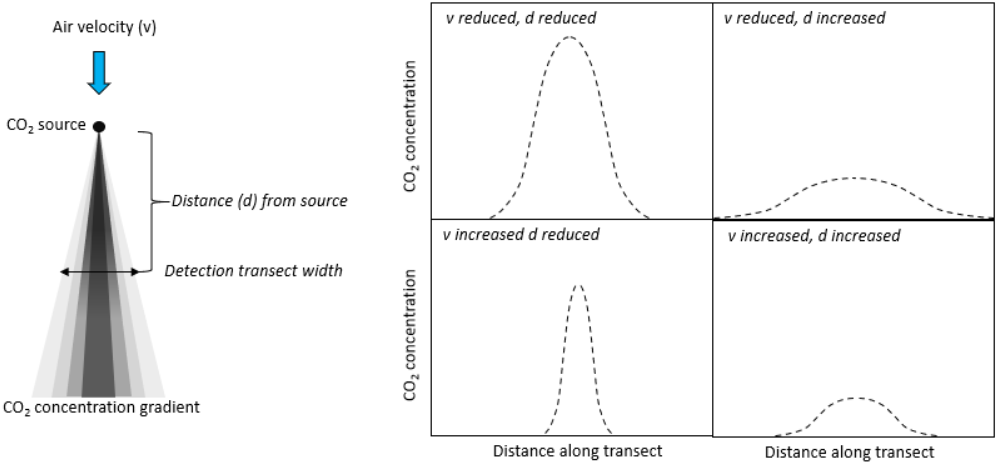
Expected changes in CO_2_ concentration in relation to sensor distance from the source and wind speed. The distribution of CO_2_ emitted at a constant rate from a point source in space is roughly expected to follow an increasing radius of the iso-concentrations as the gas diffuses out, modulated by air flow, which will tend to distribute the CO_2_ downwind of the source, with distance-dependent iso-concentration radii decreasing with increasing air speed.

In this work, we take a first step towards this goal by describing the use of the NoDIS method in wind tunnels behind perched captive, but unrestrained, pigeons *Columba livia domestica*, a starling *Sturnus vulgaris*, and a zebra finch *Taeniopygia guttata*. Our aims are to; (i) demonstrate detection of a CO_2_ signal downwind of unrestrained animals, and (ii) examine how this signal is affected by rate of CO_2_ emission, source-sensor distance, lateral position across the wake and wind speed. We also (iii) examine what the approach can tell us about animal state and respiratory physiology in perched pigeons, post-flight, post-handling and in response to a perceived threat. Finally, (iv) we map out future directions for the method to integrate the full CO_2_ shadow downwind of a resting animal and even behind a bird flying in a wind tunnel to derive figures for metabolic rates from animals undertaking activities that are currently assessed using conventional means, with associated limitations.

## RESULTS

### Definition of the CO_2_ wake downwind of a source

We used a defined gas mix of 4% CO_2_ /air (BOC) to carry out our calibrations in an open jet style wind tunnel custom designed for bird flight (test section width 1.8 m, length 2.2 m, height 1.5 m) in Swansea University, UK. Across a range of emission rates (0.5 L min^-1^ and 1 L min^-1^), source-sensor distances (10, 30 and 50 cm) and windspeeds (1, 5 and 10 m s^-1^), the width of the CO_2_ wake was measured using a NoDIS, LI-7500A Open Path CO_2_/H_2_O Analyzer (Lincoln, Nebraska, USA), and ranged from 5-14 cm (Fig 2). CO_2_ concentrations increased from the periphery of the wake towards a maximum in the centre and decreased with increasing distance from the source and increasing windspeed (Fig. 2). Transects across the downwind wake of a constant CO_2_ source showed that transect width increased with increasing distance from the source, apart from at the greatest windspeed of 10 m s^-1^ where the transect width remained narrow (Fig 2).

**Figure 2.**
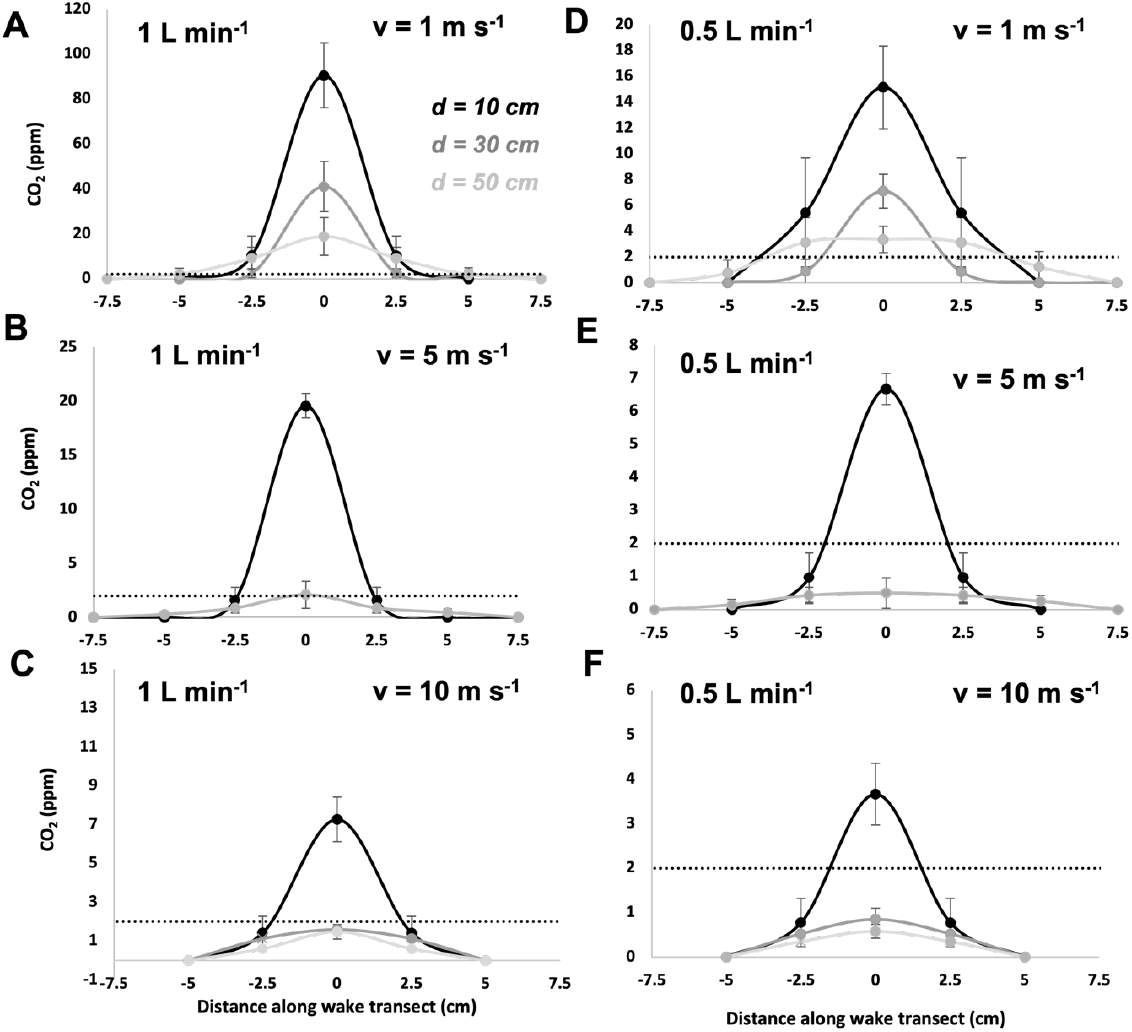
CO_2_ concentrations (mean ± S.D.) measured across the wake at different distances (d) from the source and windspeeds (v). (A-C) with a 4% CO_2_/ air mix emission rate of 1 L min^-1^. (D-F) with an emission rate of 0.5 L min^-1^. The dotted line shows the position of the 2 ppm CO_2_ concentration. N.B data were collected at -7.5, -5, 0, 2.5, 5 and 7.5 cm from the midline. At -2.5 cm means ± S.D. mirror those on the opposite side of the wake.

Using limits of detectable CO_2_ concentrations (here, 2 ppm) to define potential operational areas indicated that at an emission rate of 1 L min^-1^, the NoDIS sensor detected a clear CO_2_ signal up to 50 and 10 cm from the source at wind speeds of 1 and 10 m s^-1^, respectively (Fig. 2A-C). The same was found when the emission rate was halved (0.5 L min^-1^) (Fig. 2D-F).

### Bird CO_2_ wake exhalation signatures

In a closed system wind tunnel at the Max Planck Institute for Biological Intelligence, Germany, we positioned the sensor 46 cm behind a zebra finch, starling and pigeon with the windspeed of set to 2 m s^-1^. Clear and regular peaks in CO_2_ were detectable for all three (Fig. 3). These signals allow calculation of breathing frequency and the integral under the signal. However, as indicated by the calibration work, the quality of the signal depends on the rate of CO_2_ emission, and smaller species had less consistency in their peaks (Fig. 3). The signal was clearest when the infrared path was aligned with the tail and body (as opposed to the head) indicating that the exhaled CO_2_ attached to the body. Some variation in the signal amplitude is expected due to movement of the head, which would influence integral calculations, but birds moved their heads less with the tunnel air turned on. Measures of breathing frequency should not be affected by head movement unless an exhalation is directed away from the sensor path.

**Figure 3.**
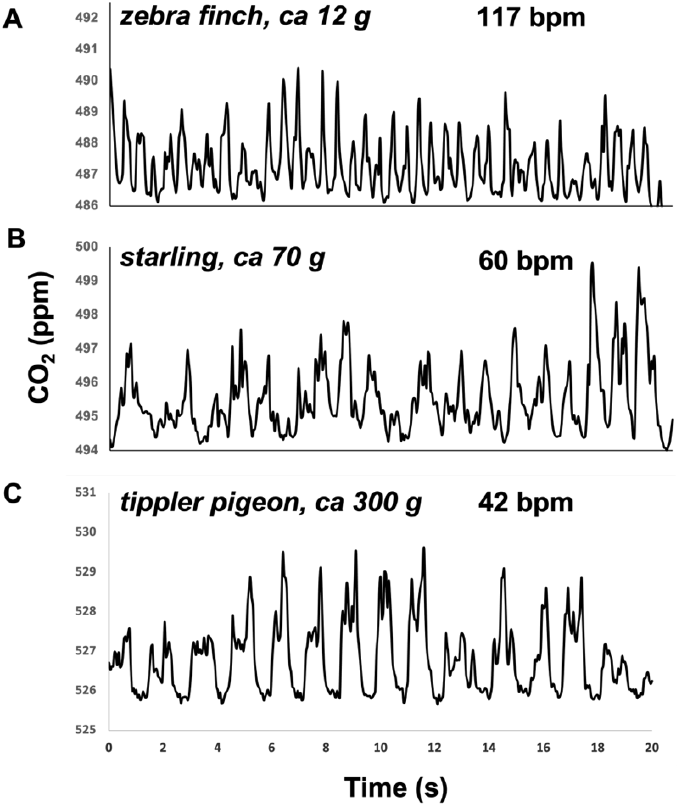
Raw CO_2_ exhalation signatures of three bird species of different body mass. In rested pigeons, CO_2_ signatures were typically M-shaped (Fig 3C and Fig 4), with the second of the two peaks often being the greatest. Here, the rate of change in CO_2_ was typically greater leading up to the second CO_2_ peak, compared to the first, and lowest during the decline following the second peak. In rested pigeons, there were short plateaus in CO_2_ concentration between exhalations that were equal to baseline measurements. In contrast, immediately post handling, and post exercise, or in smaller birds, the waveform had only a single peak and lacked plateaus between exhalations and concentration minima exceeded background CO_2_ concentrations.

### Animal state and respiratory physiology

In tippler and homing pigeons, we observed within-individual responses to different treatments. Respiration rates and, in most instances, CO_2_ production, increased during the period of exposure to a stuffed buzzard relative to a rested state, while either no response, or a smaller response, was observed when presented with a control novel object, a doll (e.g., Fig. 5, see tables S1 and S2 for statistical results for homing and tippler pigeons, respectively).

**Figure 4.**
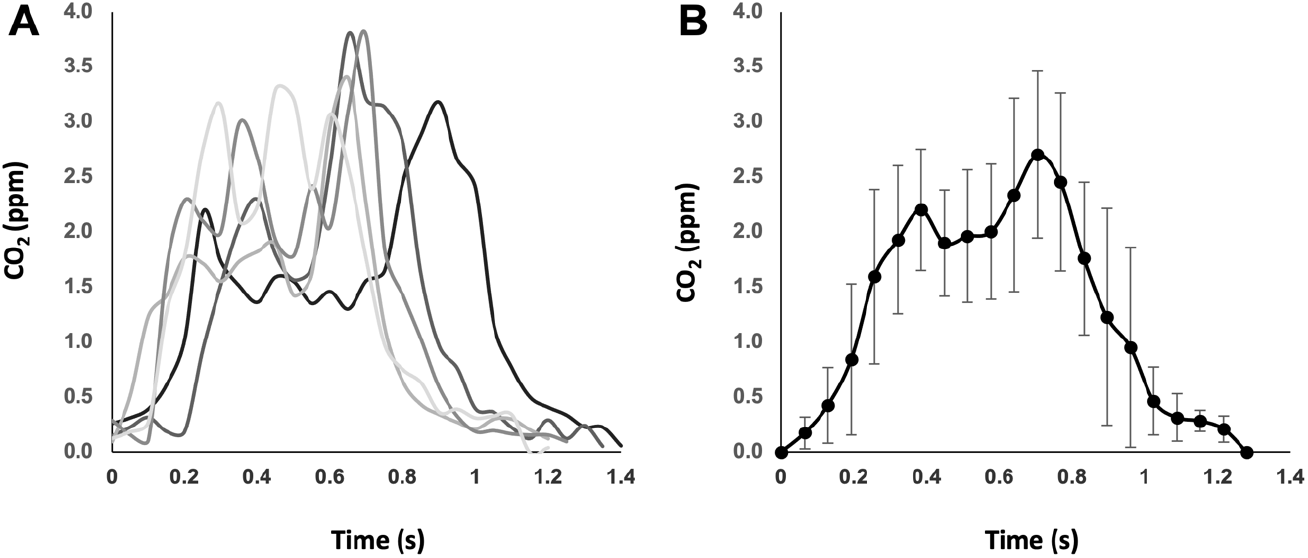
CO_2_ concentration over time for 5 consecutive breaths in a tippler pigeon stationary on a perch in a wind tunnel with an air speed of 7 m s^-1^. A) raw data sampled at 20 Hz. B) Means ± S.D every 20^th^ of a breath.

**Figure 5.**
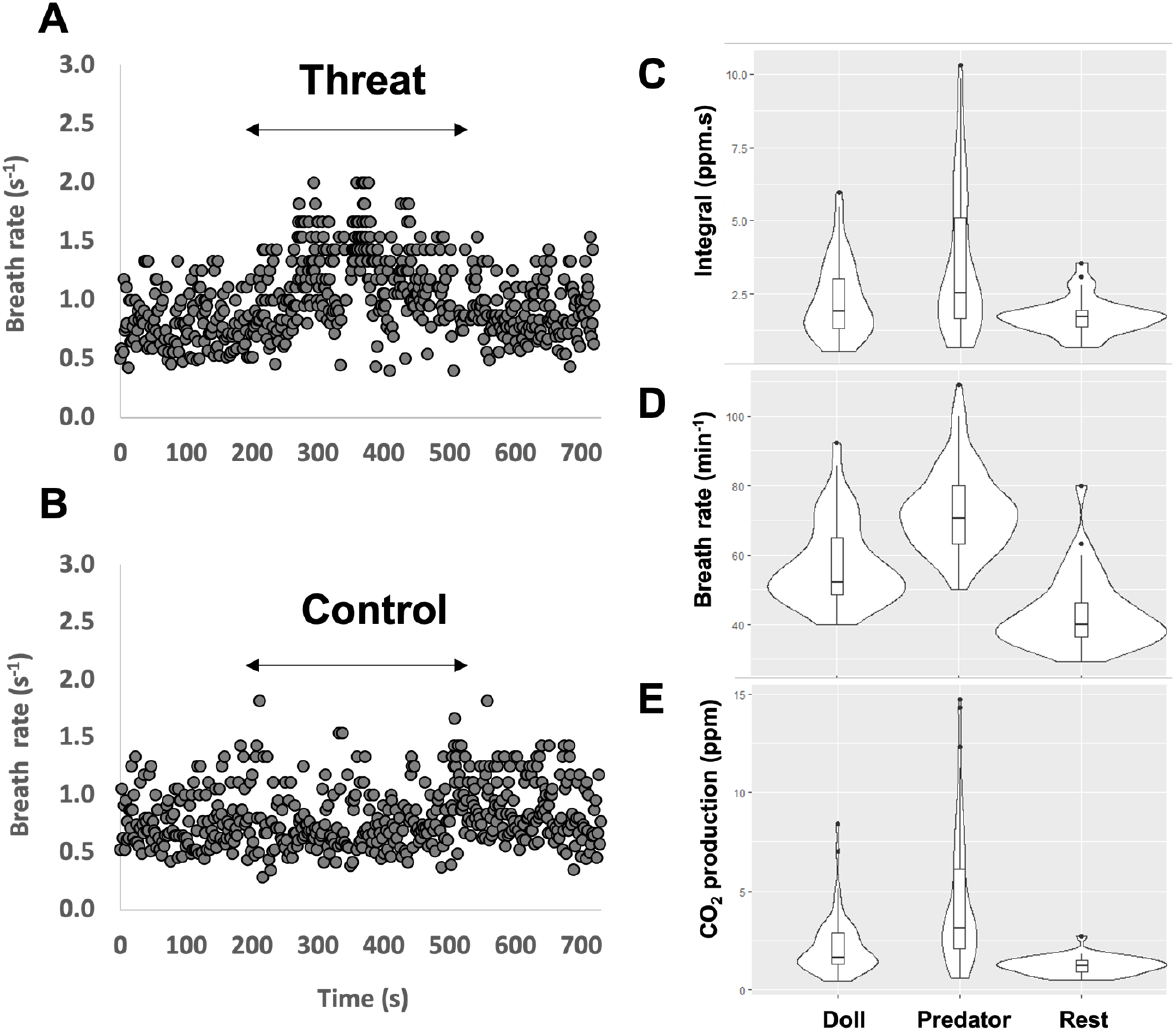
A homing pigeon’s breathing parameters in response to a perceived threat, control novel object and at rest. (A) changes in respiration rate of a rested individual exposed to a stuffed buzzard and (B) a control novel object (doll), where data points represent single breaths and arrows indicate the duration of exposure to the stimulus. Associated changes in the (C) integral of each breath, D) breath rate and E) CO_2_ production (the product of the breath rate and integral) for 1 min of data per condition (see tables S1 and S2 for statistical results).

Immediately after handling, maximum breathing rates in tippler pigeons ranged from 1.5 - 3.3 breaths s^-1^. This decreased to minimum breathing rated ranging 0.28 - 1 breaths s^-1^ within 1 minute, with most of the decline occurring within the first 10 s. Figures 6A-D show example post handling data from a homing pigeon. The integral under the exhalation peaks increased over time, although this was highly variable (Fig. 6B), and CO_2_ production decreased (Fig 6C). A negative curvilinear relationship was observed between the total CO_2_ concentration measured per breath and breath rate (Fig 6D). Figure 6E shows inter-individual variation in breath rate over time post-handling.

**Figure 6.**
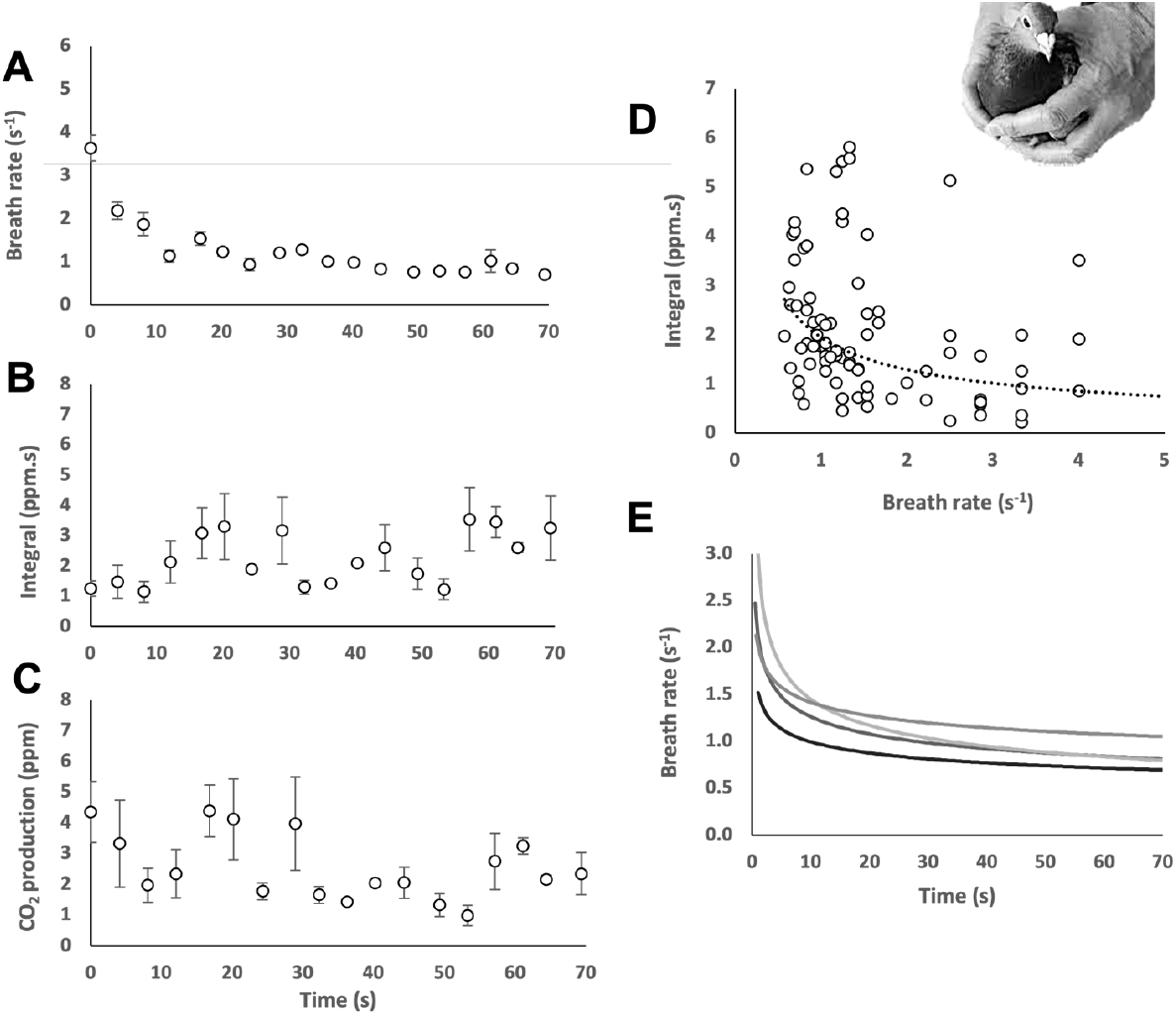
Recovery post handling and introduction to the tunnel. **A)** Respiration rate decreased to resting values within one minute, **B)** The integral increased over that minute, **C)** CO_2_ production decreased over a minute. Data points for A-C are means ± S.E over 4 seconds for a single tippler pigeon. **D)** Total CO_2_ concentration measured per breath increased with decreasing breath rate. **E)** Individual variation in breath rate over time. Each line is the best fit for an individual tippler pigeon.

In an example of post-flight recovery in a homing pigeon, breath rate declined from a maximum of 6.7 breaths s^-1^ to a minimum of 0.6 breaths s^-1^ within 1 minute (Fig. 7A). The integral of the exhalation peaks increased gradually with recovery time (Fig. 7B), whereas CO_2_ production decreased rapidly (within 10 s) (Fig. 7C). In all examples, there was a negative correlation between the integral and breath rate post flight (Fig. 6D).

**Figure 7.**
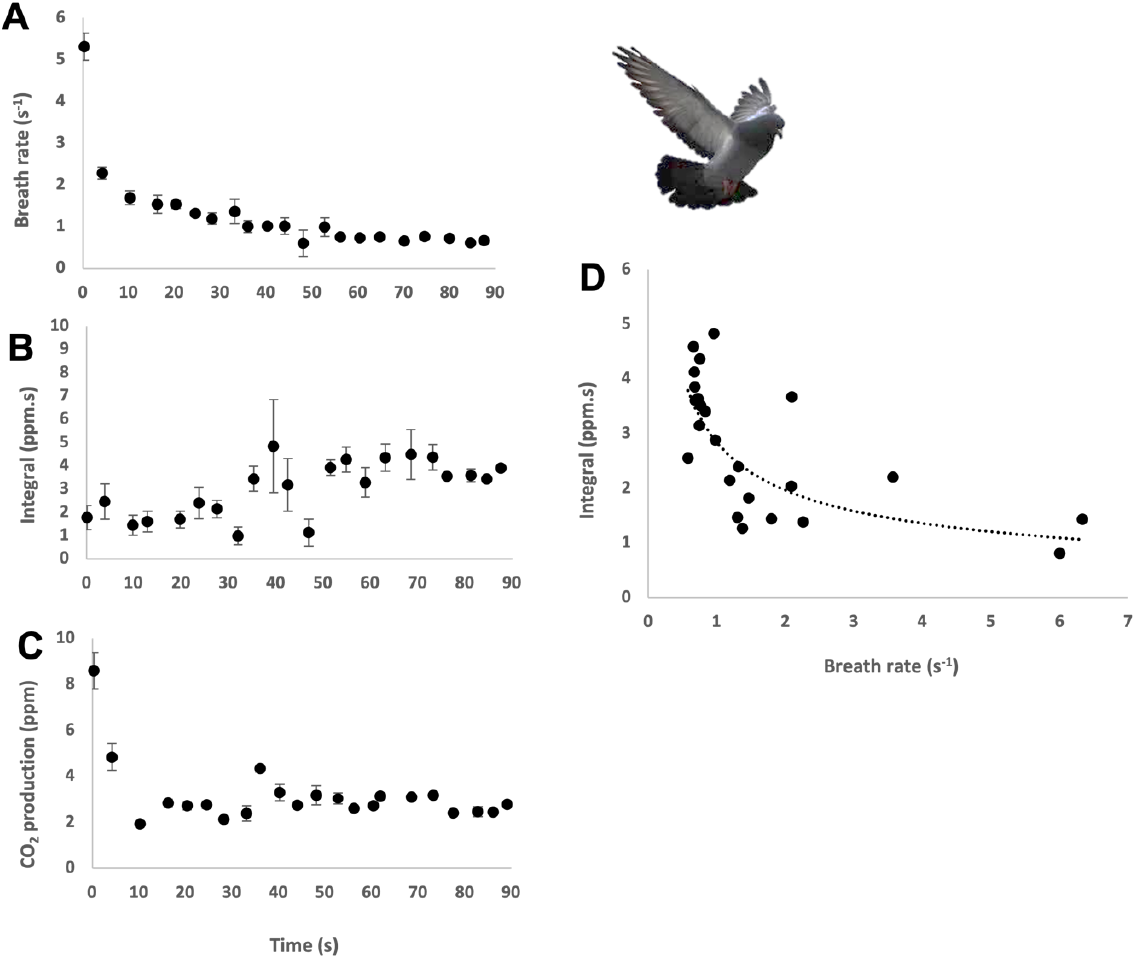
Recovery of a homing pigeon after 10 minutes of flight at 10 m s^-1^. **(A)** breath rate (s^-1^), (**B)** integral under the exhalation peak (ppm.s), (**C)** CO_2_ production (ppm), (**D)** integral versus breath rate. Datapoints represent a mean over 4 seconds and error bars in A-C are S.E..

## DISCUSSION

There is abundant literature on how amniote breathing frequency, together with tidal volume, modulates metabolic rate (Hallam & Dawson 1993; Kohin, Williams & Ortiz 1999); on its uses in the measurement of stress responses (Greenacre & Lusby 2004; Fucikova *et al*. 2009; Torné-Noguera, Pagani-Núñez & Senar 2013; Doss & Mans 2016; Doss & Mans 2017; Liang *et al*. 2018); its involvement in temperature regulation ((El Hadi & Sykes 1982; Brent *et al*. 1984; Bucher & Bartholomew 1986); entering, and arousing from, torpor (Withers 1977); and how it scales with body mass (Frappell & Baudinette 1995; Frappell, Hinds & Boggs 2001; Mortola & Seguin 2009). Here, we demonstrate that a NoDIS system can be used to quantify real-time changes in respiration rate with breath-by-breath resolution of CO_2_ concentration when the sensor is positioned in the wake of an animal, rather than integrated into a mask or alternative system that requires restraint or tethering (Butler, West & Jones 1977; Franz & Goller 2003; Wilson *et al*. 2019). In fact the system is so sensitive that, for birds as large as pigeons, two sub-peaks in CO_2_ were evident within each exhalation signature, suggesting the anterior and posterior air sacs of the respiratory system empty slightly out of phase with one another (Bretz & Schmidt-Nielsen 1971; Maina 2005; Perry, Lambertz & Shmitz 2019). Furthermore, by multiplying the frequency by the integral under the CO_2_ exhalation signature to provide a proxy for CO_2_ production, we were able to document responses to, and recovery from, stressors or exercise over fine-scales (cf (Franz & Goller 2003)).

### Limitations in detection of CO_2_ downwind of a constant CO_2_ source

The performance of the system depends on there being discernible pulses in CO_2_, and here we describe the operational limits for this. The NoDIS system that we used is reported to have an RMS noise of 0.16 ppm, which we could confirm with our baseline measurements. The variability that we obtained in our CO_2_ signals in our calibration trials in the wind tunnel (Fig. 2) was due to inconsistencies in both the rate of expulsion of the CO_2_ and in the air flow (although the wind tunnel had a high degree of laminar flow) and increased with the magnitude of the signal.

As predicted, varying wind speed affected the measured CO_2_ concentrations according to the position of the NoDIS sensor relative to the gas source and the rate of gas emission. The detection limits of the sensor (Fig. 2) indicate that, for CO_2_ emission rates of 1 L min^-1^, measurements can be made up to 50 cm immediately downwind of the source if the wind speed is 1 m s^-1^ but this reduces to 10 cm at wind speeds 10 m s^-1^. The detectable CO_2_ wake transect widths in these instances are 10 and 6 cm, respectively, indicating angles of 5.71 and 3.43° either side of a perpendicular line from the source to the sensor. With half the emission rate (0.5 L min^-1^), transect widths become 5 and 4 cm and angles become 14.03 and 11.31°.

These calibration figures provide broad working limits for researchers wishing to work with animals the size of the birds used in our study and go some way to helping define when the system might operate for different study animals and conditions. However, beyond these calibrations, further variability in the CO_2_ signal will occur due to; (i) movement of the animal (effectively displacing the CO_2_ source relative to the NoDIS sensor), (ii) the pulsed nature of CO_2_ expiration during breathing, and (iii) as a result of the air flow variability itself (direction and speed) if the system is used outside.

### Limitations in detection of CO_2_ downwind of an animal-based CO_2_ source

Animal movement is a major factor in modulating the CO_2_ signal and we suggest that this can be broadly broken down into three categories; whole body movement (where the animal moves from one site to another), body rotation (where the body turns about its vertical axis only) and head movement only. Ideally, a means to capture a full transect of the wake is required to minimise these sources of variation, whether that be achieved by moving the sensor closer to the study animal or working with a greater NODIS emitter distance or more sensors. In all cases, we recommend that researchers using NoDIS systems film their study animal if possible so that the role of movement can be ascertained. Clearly, for an animal that moves laterally, such as a bird on a perch, thereby displacing the CO_2_ source, this can result in the sensor operating outside the detection plume. In fact, such animal movement would preclude our approach for many animals for much of their time.

Complementary to the methodology demonstrated here, however, there are situations where animals may remain immobile, or at least stay in one spot for extended periods. For example from incubating (Gabrielsen *et al*. 1991), sleeping (Tillmann 2009) or torpid (Nowack, Stawski & Geiser 2017) animals, to basking reptiles (Mukherjee et al 2017), to birds such as flycatchers (Muscicapidae) and raptors which may regularly use particular look-out posts as vantage points (Fitzpatrick 1980), and territorial birds singing (Odom *et al*. 2014). Body rotation is expected to change the CO_2_ signal in a predictable manner because it effectively results in either a lateral displacement of the CO_2_ source (for example if an incubating bird facing upwind rotates 90°) and/or changes the distance between source and sensor (if the bird rotates 180°), both of which have nominally predictable effects on the concentration of the CO_2_ reaching the sensor if properly calibrated. Head movement, especially in long-necked study animals such as swans, can obviously lead to appreciable CO_2_ source displacement and the extent to which this changes the signal will depend greatly on the situation. A resting swan, for example, is predicted to have an expiration emission rate that is some 7.5 times that of a pigeon (Frappell & Baudinette 1995) which may make an immediately downwind, but distanced, NoDIS sensor, less sensitive per cm degree of lateral movement (see the extent of the flat top to the distribution in Fig. 2) than it would be in a pigeon although, of course, the pigeon would move its head absolutely less.

The carefully controlled wind conditions of the wind tunnel standardize an important element of the protocol and the value of the CO_2_ concentration data over time in the wild will be critically dependent on the variability in wind speed and the turbulence. Variation in wind speed (gustiness) increases with overall mean wind speed so the CO_2_ pulses detected by the NoDIS system will vary accordingly, specifically having the period of the exhalent pulse contracted or expanded, with accompanying changes in pulse height. Overall, this should not change respiration rate values measured over a number of cycles, but it will alter interpretation of patterns of air exhalation (see below) and may alter the values of the integrals of the CO_2_ concentration under the expiration pulse (see below), although expanded exhalation periods should be accompanied by decreased CO_2_ production. These issues may be largely mitigated by having high resolution measurement of wind speed (and vector) at the site so that, if necessary, corrections could be applied or at least data filtered to exclude aberrant gusts or periods of calm.

Another option, for specific cases where there is no wind, is to blow air at an appropriate rate past the study animal. This is easiest for animals in prescribed hollows, such as bats in their roosting boxes but may also work for animals outside. The noise, general disturbance and potential for wind effects on metabolic rate may severely limit this option though.

### Wake respirometry: Future developments

Given the accuracy of the NoDIS system in measuring CO_2_ concentrations in precise locations downwind of the study animal, it is tantalising to speculate whether this approach might enable researchers to determine metabolic rate in free-living animals. Effectively, the integral of every CO_2_ pulse should be proportional to the sum of the CO_2_ expired in that exhalation, and should indicate relative changes in metabolic rate. It could therefore be used to examine recovery in animals that have engaged in movement outside the sample area before returning, in a manner similar to our pigeons (Fig. 7). Birds flying back to their nest would be an obvious example.

Derivation of absolute metabolic rate is more challenging. However, data from multiple NoDIS sensors within the CO_2_ footprint could be integrated and summed to provide the complete CO_2_ footprint per exhalation and over longer time periods. Alternatively, a series of inhalant tubes could sample the wake and mix the air for analysis by a single NoDIS sensor to attempt derivation of total CO_2_ emission. Either way, some sort of calibration would be desirable but simulation of CO_2_ emissions from the study animal on site *post hoc* by bleeding gas from models may help.

Overall, this work has demonstrated that the new, portable generation of CO_2_ sensors can provide insight into stress and respiratory patterns in unrestrained animals, and as a result, that they could be used to document stress responses in wild animals. With suitable consideration of the limitations imposed by factors such as animal movement and wind variability, wake respirometry should have a future that will help with a diverse suite of issues, such as determination of the extent to which urbanisation (Charmantier *et al*. 2017), tourism (Mullner, Linsenmair & Wikelski 2004), or natural disasters (Nowack, Stawski & Geiser 2017) might affect target animals, and indeed, comparison of the effects of different stressors (Clinchy, Sheriff & Zanette 2013). Beyond that, it also opens the way for potential measurements of the metabolic rate of unrestrained birds, both resting, and in the case of wind tunnels, in flight.

## STAR METHODS

### Definition of the CO_2_ shadow downwind of a source

#### Apparatus

Non-dispersive infrared spectroscopy uses infrared radiation that is emitted across an (open) path of defined length, across which the CO_2_ is to be measured, with the radiation being detected at the distal end of the path by a lead selenide sensor. Both water vapour and CO_2_ absorb the radiation so gas densities can be determined by considering the absorption with respect to a reference. We used the LI-7500A Open Path CO_2_/H_2_O Analyzer (Lincoln, Nebraska, USA), which has an emitter-sensor distance of 125 mm, resolution 0.01 ppm, and error within 1% of reading. During our use of this system, we deployed it in a vertical orientation and set it to sample at 20 Hz (RMS noise 0.16 ppm at 370 ppm CO_2_).

#### Calibrations

We assessed the viability of our approach by conducting trials under varying conditions. We used a defined gas mix of 4% CO_2_ /air (BOC) to carry out our calibrations in an open jet style wind tunnel custom designed for bird flight (test section width 1.8 m, length 2.2 m, height 1.5 m) in Swansea University, UK. Rubber tubing (10 mm outer diameter) and a variable flow metre were used to release the gas mix at two fixed flow rates (0.5 and 1 L min^-1^), via a metal tube inserted through the ceiling of the tunnel, which extended to a central position 60 cm inside the test section. The NoDIS sensor was positioned on a stand at the same height as the source to record CO_2_ (ppm) at three source-sensor distances (10, 30, 50 cm) and three wind speeds (1, 5 and 10 m s^-1^). At each emission rate, windspeed, and distance (d) from the source, ten seconds of CO_2_ (ppm) data was logged (20 Hz) every 5 or 2.5 cm along the wake transect until the signal was no longer detectable.

### CO_2_ signals from captive birds

#### Captive birds used in wind tunnel trials

Data were recorded from adult captive tippler pigeons (n=4, *ca*. 300 g), homing pigeons (n=5, *ca*. 400 g), a starling (*ca*. 70 g) and a zebra finch (*ca*. 10 g) in a closed system wind tunnel at the Max Planck Institute for Biological Intelligence, Germany. Birds were kept in aviaries beside a closed system wind tunnel according to §11 Permission (§11 TierSchG) and were accustomed to being inside the tunnel for flight training. Animal experiments performed in Seewiesen, Germany, were conducted according to the regulations of the government of Upper Bavaria (Germany protocol numbers: AZ 55.2-1-54-2532-86-2015; 311.5-5682.1/1-2014-021). At Swansea University, data were recorded from a hand-reared homing pigeon (female, *ca*. 350 g) accustomed to flight training in the open jet wind tunnel. Here, homing pigeons were housed in an outdoor loft with aviary under an establishment licence. Experiments were carried out under the project licence (X5770C662) and ethical permission for this work was given by Swansea University AWERB (200418/65).

#### Measurements from birds in different physiological states in wind tunnels

We investigated whether our set-up could detect responses of the tippler and homing pigeons to the following treatments; i) release after being handled, which was assessed by keeping birds in a darkened box before they were held by an experimenter for 2 minutes and then introduced to the tunnel perch, and proximity to ii) a potential threat (stuffed buzzard *Buteo buteo*) and iii) a similar-sized control novel object (rag doll). Within-individual responses to the buzzard and doll were assessed when the birds were in a rested state on the perch.

The perch (55 cm tall, 8 cm wide) was positioned in the centre of the test section while the NoDIS was fixed to a stand positioned 46 cm downstream. The perch width was relatively small to prevent birds moving to the left or right relative to the sensor. Birds always faced into the wind when the tunnel was on. Fixed windspeeds (7 and 10 m s^-1^ for tippler and homing pigeons, respectively) were chosen to ensure a clear signal with minimal variation in CO_2_ concentration between exhalations.

The wind tunnel room was 17-19°C. Lights in the study room were dimmed to a low level and noise additional to that of the wind tunnel was kept to a minimum. Two experimenters were in the room during all trials, which were conducted during normal active hours. A typical trial consisted of a bird being placed inside a darkened carrier box (dimensions 50 × 35 × 30 cm) for 10 minutes in the wind tunnel room. The bird was then removed from the box and held for 2 minutes with fingers around both sides of the body. The same experimenter restrained all birds (Rabdeau *et al*. 2019). At the same time, a baseline CO_2_ trace was recorded in the absence of the bird with the tunnel on. The bird was then placed on the perch upstream of the sensor and left to sit quietly. After 20 min, either the taxidermy buzzard or rag doll was presented outside the test section and upstream of the study bird and held there for 2 minutes. Another 20 minutes of quiet time followed before the second stimulus was presented. The order in which the stimuli were introduced within trials was randomised and birds were presented with each stimulus only once to avoid habituation.

For a comparison of the CO_2_ exhalation signal between birds of different body size at a single wind speed and bird-sensor distance (46 cm and 2 m s^-1^), resting data were collected from a perched tippler pigeon, starling, and zebra finch in the Max Planck wind tunnel. Twelve starlings were flown together at 10 m s^-1^ for 10 minutes as part of their usual training regime, after which, one individual was kept within the tunnel to recover on a perch. Similarly, ten zebra finches were flown together at 8 m s^-1^ for 10 minutes, and one individual was kept within the tunnel for breath recordings.

Data were also collected behind a homing pigeon in the test section of the tunnel at Swansea University after 11 minutes of flight training at 10 m s^-1^ for post-flight recovery data. The bird was perched 20 cm upstream of the NoDIS (perch height 60 cm) and the wind speed remained at 10 m s^-1^ during data collection after the flight training. Lights were dimmed at the end of a training session for respirometry measurement and experimental temperatures ranged from 18.8-19.2°C.

### Data processing and extraction

CO_2_ data (ppm) were corrected for baseline drift in OriginLab 2021 using linear interpolation between baseline data collected in the absence of a bird at the beginning and end of trials or at regular intervals throughout calibration experiments in the absence of CO_2_ emission. Calibration measurements, exhalation and breath cycle parameters were examined using in house software (DDMT, Wildbyte Technologies, http://wildbytetechnologies.com). Changes in CO_2_ over time were isolated for each expiration from drift-corrected and baseline-corrected data. Breathing frequency (breaths per unit time) was calculated as the reciprocal of the breath cycle period. A proxy for CO_2_ production was calculated by multiplying breathing frequency by the integral of the CO_2_ signal.

### Statistical analyses

Statistical analyses were conducted in R Studio using R version 4.0.3 (R Development Core Team 2019). Non-parametric Kruskal-Wallis and Dunn (holm adjusted) post-hoc tests were conducted to investigate within-individual differences in pigeon breath rate, integral of the CO_2_ signature and CO_2_ production (breath rate x integral) under three different conditions: in a rested and undisturbed state; when exposed to a control novel object (rag doll); and when exposed to a potential threat (stuffed buzzard). In all cases, assumptions for parametric one-way ANOVAs were not met and this was confirmed by examining qq plots and histograms of model standardised residuals as well as Shapiro Wilk tests to confirm a distribution significantly different from normal. For each pigeon, one minute of breath-by-breath data was used per condition. Data were investigated separately for each pigeon because of the variability of signal strength that is expected due to each bird’s varying body form and head movement, our small sample size, and additional variation in measurements expected due to individual differences in body mass.

---

**Table S1.** Results of Kruskal-Wallis tests and Dunn (holm adjusted) post hoc tests to investigate differences in the respiratory parameters of individual homing pigeons exposed to different stimuli.

| ID | Parameter | Median | $\epsilon^2$ | $X^2$ (df) | P | n | Comparison | z | P.unaj | P.adj |
| --- | --- | --- | --- | --- | --- | --- | --- | --- | --- | --- |
| HP1 | Integral (ppm.s) | Rest: 1.72<br>Doll: 1.92<br>Predator: 2.52 | 0.089 | 16.71 (2) | **<u>0.0002</u>** | 169 | doll - predator<br>doll - rest<br>predator - rest | -1.47<br>0.70<br>2.43 | 0.142<br>0.486<br>0.015 | 0.283<br>0.486<br>**<u>0.046</u>** |
|  | Breath rate (/min) | Rest: 40<br>Doll: 52.17<br>Predator: 70.59 | 0.555 | 94.19 (2) | **<u><0.001</u>** | 169 | doll - predator<br>doll - rest<br>predator - rest | -5.62<br>4.09<br>9.44 | <0.001<br><0.001<br><0.001 | **<u><0.001</u>**<br>**<u><0.001</u>**<br>**<u><0.001</u>** |
|  | CO<sub>2</sub> production (ppm) | Rest: 1.24<br>Doll: 1.65<br>Predator: 3.14 | 0.310 | 53.49 (2) | **<u><0.001</u>** | 169 | doll - predator<br>doll - rest<br>predator - rest | -4.58<br>1.73<br>7.10 | <0.001<br>0.083<br><0.001 | **<u><0.001</u>**<br>0.083<br>**<u><0.001</u>** |
| HP2 | Integral (ppm.s) | Rest: 1.71<br>Doll: 1.47<br>Predator: 1.94 | n/a | 3.26 (2) | 0.196 | 219 | n/a | n/a | n/a | n/a |
|  | Breath rate (/min) | Rest: 40<br>Doll: 50<br>Predator: 63 | 0.183 | 41.52 (2) | **<u><0.001</u>** | 219 | doll - predator<br>doll - rest<br>predator - rest | -6.44<br>-3.84<br>3.42 | <0.001<br><0.001<br><0.001 | **<u><0.001</u>**<br>**<u><0.001</u>**<br>**<u><0.001</u>** |
|  | CO<sub>2</sub> production (ppm) | Rest: 1.76<br>Doll: 1.23<br>Predator: 2.00 | 0.037 | 10.06 (2) | **<u>0.007</u>** | 219 | doll - predator<br>doll - rest<br>predator - rest | -2.74<br>-2.88<br>0.18 | 0.006<br>0.004<br>0.854 | **<u>0.012</u>**<br>**<u>0.011</u>**<br>0.854 |
| HP3 | Integral (ppm.s) | Rest: 2.18<br>Doll: 2.80<br>Predator: 2.39 | 0.035 | 8.38 (2) | **<u>0.016</u>** | 184 | doll - predator<br>doll - rest<br>predator - rest | 2.57<br>2.66<br>-0.10 | 0.010<br>0.007<br>0.918 | **<u>0.020</u>**<br>**<u>0.023</u>**<br>0.918 |
|  | Breath rate (/min) | Rest: 41.38<br>Doll: 40<br>Predator: 57.14 | 0.440 | 81.61 (2) | **<u><0.001</u>** | 184 | doll - predator<br>doll - rest<br>predator - rest | -7.84<br>-1.53<br>7.74 | <0.001<br>0.126<br><0.001 | **<u><0.001</u>**<br>0.126<br>**<u><0.001</u>** |
|  | CO<sub>2</sub> production (ppm) | Rest: 1.53<br>Doll: 1.88<br>Predator: 1.95 | 0.045 | 10.22 (2) | **<u>0.006</u>** | 184 | doll - predator<br>doll - rest<br>predator - rest | -0.78<br>1.89<br>3.08 | 0.435<br>0.058<br>0.002 | 0.435<br>0.117<br>**<u>0.006</u>** |
| HP4 | Integral (ppm.s) | Rest: 2.67<br>Doll: 3.46<br>Predator: 3.98 | 0.054 | 11.27 (2) | **<u>0.003</u>** | 174 | doll - predator<br>doll - rest<br>predator - rest | -1.43<br>1.72<br>3.32 | 0.154<br>0.087<br><0.001 | 0.154<br>0.173<br>**<u>0.003</u>** |
|  | Breath rate (/min) | Rest: 41.28<br>Doll: 46.15<br>Predator: 46.15 | 0.195 | 35.30 (2) | **<u><0.001</u>** | 174 | doll - predator<br>doll - rest<br>predator - rest | 0.98<br>5.40<br>4.30 | 0.033<br><0.001<br><0.001 | 0.033<br>**<u><0.001</u>**<br>**<u><0.001</u>** |
|  | CO<sub>2</sub> production (ppm) | Rest: 1.75<br>Doll: 2.60<br>Predator: 2.76 | 0.140 | 25.86 (2) | **<u><0.001</u>** | 174 | doll - predator<br>doll - rest<br>predator - rest | -0.78<br>3.72<br>4.60 | 0.435<br><0.001<br><0.001 | 0.435<br>**<u><0.001</u>**<br>**<u><0.001</u>** |

**Table S2.**
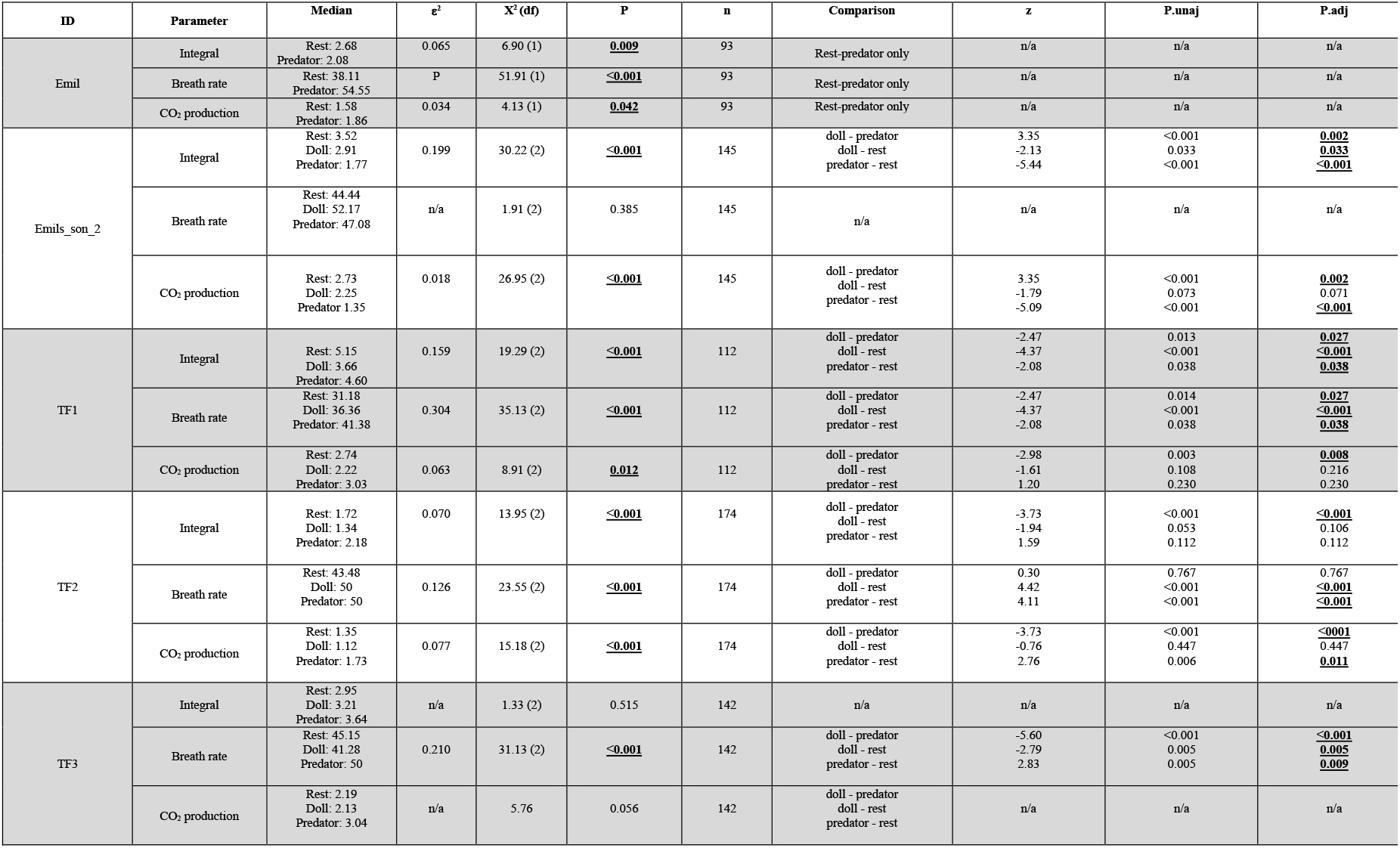
Results of Kruskal-Wallis tests and Dunn (holm adjusted) post hoc tests to investigate differences in the respiratory parameters of individual tippler pigeons exposed to different stimuli.

## ACKNOWLEDGEMENTS

This work was supported by the European Research Council under the European Union’s Horizon 2020 research and innovation program (starting grant 715874 to ELCS) and a Max Planck Sabbatical Fellowship (to ELCS).

## AUTHOR CONTRIBUTIONS

Conceptualization, ELCS, RPW, KARR; Methodology, ELCS, RPW, KARR; Investigation, KARR, HR, CR; Resources, MW, Formal Analysis, KARR; Writing – Original Draft, ELCS, RPW, KARR; Writing – Review & Editing, ELCS, RPW, KARR, MW, HR, CR; Project Administration, ELCS, KARR; Funding Acquisition, ELCS.;

## Notes

### Competing Interest Statement

The authors have declared no competing interest.

